# Predicting alcohol consumption from resting-state functional connectivity: Feature relevance and the potential of meta-matching

**DOI:** 10.64898/2026.10.19.759632

**Authors:** Marco Bottino, Michael N. Smolka, Justin Böhmer, B.T. Thomas Yeo, Patrick Bach, Falk Kiefer, Bernd Lenz, Andreas Heinz, Gianna Spitta, Annika Rosenthal, Kerstin Ritter, Roshan Prakash Rane, Shreyash Garg, Emanuel Schwarz, ReCoDe Consortium, Henrik Walter, Michael Marxen

**Author notes:** shared senior authorship (Henrik Walter, Michael Marxen). **Corresponding author:** Michael Marxen, Ph.D., Section of Systems Neuroscience, Department of Psychiatry and Psychotherapy,Technische Universität Dresden, Würzburger Straße 35, 01187 Dresden, Germany. Other authors emails and ORCID ID Marco Bottino Michael N. Smolka Justin Böhmer B. T. Thomas Yeo Patrick Bach Falk Kiefer Bernd Lenz Andreas Heinz Gianna Spitta Annika Rosenthal Kerstin Ritter Roshan Prakash Rane Shreyash Garg Emanuel Schwarz Henrik Walter.

## Abstract

Alcohol use disorder (AUD) is among the most prevalent mental disorders. Associations between resting-state fMRI functional connectivity (FC) and AUD have been reported but rarely replicated. In this preregistered study, we investigated the extent to which FC predicts alcohol consumption, as measured by the AUDIT-C score, in a large sample of non–treatment-seeking alcohol consumers (N = 739), including 528 participants with varying degrees of AUD severity. To optimize performance, we compared kernel ridge regression (KRR) trained directly on this sample with transfer-learning models based on multilayer meta-matching embeddings, from which we derived data-driven relevance maps at the connection, region, and network levels. We tested split-half reliability, reproducibility across cohorts, and outcome specificity of the relevance maps. Both models predicted AUDIT-C scores moderately but significantly (*r* ≈ 0.15). Meta-matching did not outperform KRR in prediction. Relevance maps showed limited split-half reliability, consistent with the modest predictive performance, but reproduced moderately well across two independent cohorts. These maps indicate reduced connectivity among limbic regions, the anterior middle temporal gyrus, the dorsomedial prefrontal cortex, and dorsal attention network regions. In contrast, hypothesized hyperconnectivity patterns involving the striatum, anterior cingulate cortex, and precuneus were less reliable and mostly non-significant. Overall, our findings suggest that resting-state FC contains reproducible but weak signals related to alcohol consumption. Transfer learning improved relevance-map reliability but not prediction performance and reduced outcome specificity. These results support the utility of interpretable machine learning for identifying connectivity patterns related to drinking behavior, while also highlighting current limitations in its predictive power.

## Introduction

Alcohol Use Disorder (AUD) is a highly prevalent psychiatric condition marked by compulsive alcohol consumption that potentially leads to adverse personal, social, and health-related impairments, significantly contributing to the global burden of disease (Shield et al., 2020). Alcohol consumption is among the leading global risk factors for mortality (Collaborators, 2018).

Functional magnetic resonance imaging (fMRI), which measures brain activity via the blood oxygen level-dependent (BOLD) signal, is widely used to study alterations in brain function in psychiatric conditions, including AUD (Bottino et al., 2025; Rodriguez et al., 2025). In particular, the relationship between resting-state functional connectivity (rsFC) and AUD has been widely explored in the literature, mostly by comparing groups of subjects with diagnosed AUD with healthy controls (HC). In our previous work, we conducted a systematic review summarizing the main findings from the literature on the topic and generated literature-based “relevance maps” that identify functional connections, regions, and networks most consistently associated with AUD (Bottino et al., 2025). The literature has particularly implicated alterations within– and between-network connectivity involving the salience network (SN), default mode network (DMN), and executive control network (ECN), which are central to the triple-network model of psychopathology (Elsayed et al., 2024; Guerrero et al., 2025). Some of these alterations involved the reward processing system, such as increased connectivity between DMN regions, like the precuneus, and the striatum. On the other hand, the disconnection of executive prefrontal regions from the bilateral temporal poles and pre– and postcentral gyri can be associated with altered emotion regulation and inhibitory control (Deng et al., 2022; Duan et al., 2022). Overall, such alterations may help unravel the neural correlates of disorder symptoms, but our review found poor reproducibility of associations between resting-state fMRI FC and AUD, likely due to low sample sizes and methodological heterogeneity across studies.

Methodologically, multivariate machine learning techniques are increasingly used to predict psychiatric phenotypes from neuroimaging data, shifting the focus from group-level contrasts to individual-level prediction, which can help bridge the gap to clinical application. While older studies often classify individuals into discrete categories, e.g., AUD vs. HC (Li et al., 2023; Vergara et al., 2022; Zhu et al., 2018), regression-based methods offer the advantage of modeling AUD as a continuous dimension, avoiding arbitrary group thresholds. Some studies use such approaches to predict AUD severity from FC (Fede et al., 2019; Song et al., 2024); however, these efforts have typically relied on relatively small samples, limiting the robustness and generalizability of their findings. Indeed, a major challenge in behavioral regression from fMRI data is the risk of overfitting due to the high dimensionality of the input space, where the number of features often far exceeds the number of observations – even in large-scale neuroimaging cohorts (Scheinost et al., 2019; Whelan C Garavan, 2014). Small sample sizes can inflate performance estimates (Flint et al., 2021) and lead to large uncertainty in model evaluation, as reflected by large error bars and low replicability (Varoquaux, 2018).

One strategy for addressing this problem is transfer learning, in which information learned from large external datasets is leveraged to improve prediction in smaller target samples. Meta-matching is one such approach and has been shown to improve prediction in unseen datasets compared with models trained exclusively on the target sample (Chen et al., 2024; He et al., 2022). More recently, meta-matching models have demonstrated cross-dataset generalizability for cognitive phenotypes (Chopra et al., 2024), but, to our knowledge, the same has not yet been demonstrated for AUD-related outcomes.

In this paper, we used data from the ReCoDe cohort (Heinz et al., 2020; Spanagel et al., 2024) comprising non-treatment-seeking participants spanning a broad range of alcohol use and AUD severity, as well as individuals without AUD. We investigated whether the most recent multilayer meta-matching implementation, trained on five large external cohorts, improves the prediction of AUD-related variables compared to a regression model trained solely on the ReCoDe dataset. We subsequently quantified the contribution of individual functional connections to model predictions and aggregated these associations at the regional and network levels to derive data-driven relevance maps. Finally, we evaluated the reliability and outcome specificity of these maps.

Prior to accessing the ReCoDe cohort data, we conducted an exploratory methods development to optimize the prediction of alcohol consumption from FC in an independent sample drawn from the UK Biobank (UKBB; (Sudlow et al., 2015)). These analyses were used to refine the analytical pipeline and formulate the hypotheses that were subsequently preregistered before application to ReCoDe. Based on these exploratory analyses, we selected AUDIT-C, which is a brief screening measure of alcohol consumption patterns and hazardous drinking but does not directly assess dependence symptoms or alcohol-related consequences (Saunders et al., 1993), as the primary outcome, while still examining other behavioral measures associated with the disorder.

Based on our literature review and the preregistered exploratory findings, we hypothesized that meta-matching models would outperform locally trained models and yield reliable, outcome-specific relevance maps. We further hypothesized that these maps would associate the AUDIT-C score with greater between– and within-network connectivity in the limbic network and striatum, and with weaker connectivity between temporal regions and regions implicated in sensorimotor, default-mode, and cognitive-control processing, including the precuneus, parietal operculum, and left inferior frontal gyrus.

To our knowledge, this is the first study to use interpretable transfer learning on FC data in a large sample of non-treatment-seeking participants with mostly moderate levels of AUD. By identifying connectivity patterns that are both predictive at the individual level and reproducible across samples, this approach may help clarify the neural mechanisms underlying problematic alcohol use and highlight targets for future intervention studies. In the longer term, more robust and generalizable neurobiological markers could contribute to a more individualized understanding of addiction and support the development of better-targeted prevention and treatment strategies.

## Methods

### Datasets

This study focuses on data from the TRR 265 ReCoDe cohort (Heinz et al., 2020; Spanagel et al., 2024). This cohort was collected across three sites in Germany (Dresden, Berlin, Mannheim) to study underlying mechanisms and trajectories, as well as possible interventions for substance use disorders. It comprises both cross-sectional and longitudinal data on N = 1015 participants aged 18 to 65 years, including 746 participants with AUD (455 male, 291 female) and 269 controls (138 male, 131 female). AUD diagnosis was established according to DSM-5 (Association, 2013). Subjects with other mental or substance use disorders aside from AUD, tobacco use disorder (TUD), and cannabis use disorder (CUD) were excluded. This analysis focuses on cross-sectional data. The ReCoDe cohort collection procedures and the overall consortium are described in Spanagel et al. (2024). Resting-state fMRI scans were acquired from N = 846 participants in the ReCoDe cohort. Of these, N = 16 were excluded due to technical problems, and N = 42 were excluded after manual quality assurance. Of the remaining N = 788 participants, N = 49 were excluded due to missing data in the AUDIT survey, leaving N = 739 participants (AUD: N = 528) for the present analysis. We ran the preliminary analyses used to preregister the methods and hypotheses on a subset of 711 participants from the UK Biobank (UKBB) cohort, matched for gender and drinking-pattern distributions to those in the ReCoDe cohort, shortly before completing data acquisition. We provide more details on the matching process in the supplementary materials.

### MRI acquisition and preprocessing

Scanning was carried out over 3 sites on 3T MRI scanners. Functional images were acquired using a multi-band T2*-weighted gradient echo-planar imaging (EPI) sequence (500 volumes, TR = 869 ms, TE = 38 ms, 60 slices, slice thickness = 2.4 mm, voxel size 2.4 × 2.4 × 2.4 mm). We extracted several details on subject motion during acquisition: the percentage of frames with framewise displacement (FD) over 0.5 mm, the average FD as defined in Power et al. (2012), and the time courses of subjects’ motion in 6 degrees of freedom (3 translational and 3 rotational). Preprocessing of functional images was performed using SPM12 and included slice-timing correction, realignment, and fieldmap correction (TR = 698 ms, TE1 = 5.19 ms, TE2 = 7.65 ms, 64 slices, slice thickness = 2.4 mm, voxel size 2.4 × 2.4 × 2.4 mm). T1 images (TR = 2000 ms, TE = 2.01 ms, TI = 880 ms, voxel size 1 × 1 × 1 mm) were coregistered with the EPI images and normalized to MNI space. EPI images were normalized to the MNI152 standard space at a resolution of 2.4 mm using the T1-based transform. We proceeded by applying a high-pass temporal filter (Gaussian-weighted least-squares straight line fitting, with σ = 50.0 TRs). Since filtering removes the temporal mean signal, it was added back. Afterward, we performed ICA-AROMA (Pruim et al., 2015) for denoising, followed by global signal regression, to remove any remaining physiological and motion artifacts. Finally, we resliced the images to a voxel size of 2mm.

Quality assurance was performed by visually inspecting the data before and after denoising. The pre-denoising inspection identified imaging artifacts, leading to the exclusion of 38 subjects. Post-denoising inspection used carpet plots (Power, 2017) to assess residual physiological and motion-related confounds, leading to the exclusion of an additional 4 subjects.

### Functional connectivity estimation

We derived FC matrices from resting-state sequences using an atlas comprising 400 cortical and 19 subcortical regions (Schaefer et al., 2018), as used by Chen et al. (2024) and Chopra et al. (2024) since the models employed in this study were trained on FC matrices generated from this atlas (Schaefer+). Specifically, we averaged the time series of all voxels within each region, and then computed pairwise Pearson correlations across the 419 regions. Because the resulting 419×419 FC matrix is symmetric, we extracted only the lower-triangular values for each subject as input to the models, yielding 87,571 input features. These values were then Fisher z-transformed and individually standardized to have a within-subject mean of zero and a Euclidean norm of 1 across FC values, matching the input format on which the meta-matching predictors were originally trained.

### Behavioral outcome measures

The main outcome variable for quantifying alcohol consumption in this study is the AUDIT-C score, a psychometric score equal to the sum of the first 3 items of the Alcohol Use Disorder Identification Test (Saunders et al., 1993). The items concern the frequency of drinking occasions, the number of drinks per occasion, and the frequency of binge-drinking (>6 drinks per occasion). The score can therefore be interpreted as a descriptor of a subject’s consumption patterns; thus, we will use the terms AUDIT-C score and drinking patterns or alcohol consumption interchangeably in this manuscript. We conducted further exploratory analyses of other outcomes related to drinking patterns, including weekly quantity of alcohol consumed per session in grams, frequency of drinking days and binge drinking days (>60g in one session), the full AUDIT score, the sum of its items 4-10 describing disorder-related problems, frequently called in the literature AUDIT-P (Selin, 2006), the total number of DSM-5 criteria for AUD met by the participant, and the number of DSM-5 criteria met amongst the first four, associated specifically to failures in self-control (Association, 2013).

### Brain-behavior models

We compared the performance of Kernel Ridge Regression (KRR) applied directly to the ReCoDe FC data with that of two predictive models trained on embeddings derived from multilayer meta-matching. The embedding model was previously published (Chen et al., 2024) and pre-trained on five large external cohorts to predict 458 phenotypes, providing a low-dimensional representation of FC that captures shared variance across behavioral domains. When applied to the ReCoDe data, this model generates participant-specific embeddings, which were then used as input to train two downstream predictive (stacked) models. Specifically, we evaluated KRR (MMM-KRR) and Elastic Net regression (MMM-EN) as stacked models. Further methodological details on the machine learning models are available in the supplementary materials.

Following the pipeline described in Chopra et al. (2024), we trained each model using 100 repetitions of an 80–20 random train–test split. We considered sex, age, site of acquisition, and head motion (measured as mean FD) as confounders in the prediction. The effects on outcome variables of all confounders aside from site were estimated in training sets and regressed out accordingly from training and test sets, after which we applied between-subjects standardization in the same fashion, ensuring that both inputs and outcomes had zero mean and unit variance across training subjects. Similarly, we harmonized the input arrays across sites using ComBat for each split, as implemented in the neuroHarmonize Python library (Johnson et al., 2006; Yu et al., 2018), to remove site-related effects while preserving the between-site variability introduced by the confounders listed above and the target outcome. Covariate effect removal, normalization, and site harmonization were also applied to the test sets using the parameters fitted on the training set to avoid data leakage. Within each training set, we performed 5-fold cross-validation using a random search with 50 draws to tune the models’ hyperparameters.

For each run, we evaluated model performance using three metrics: the coefficient of determination (*R^2^*), mean squared error (MSE), and mean Pearson’s correlation between predictions and ground truth (*r*) across all test sets, with the latter one being our primary performance metric.

### Derivation of the data-driven relevance maps

Following Chopra et al. (2024), we computed data-driven relevance maps at three levels for each model and outcome:

1. Functional connection level: We assessed feature importance in each of the 100 80/20 splits by computing the Haufe transform, defined as the covariance between each input feature and the model’s prediction across training subjects (Haufe et al., 2014). In our case, we used the corresponding correlation coefficients instead of covariance to account for differences in variance across input features. Note that this was necessary because only the embeddings were standardized across training subjects, not the input FC features. We averaged the 100 Haufe-transformed weight arrays and reshaped them into the original 419×419 matrix format. Next, we split these weights into two matrices: a “positive” matrix representing connections positively associated with the disorder (positive weights) and a “negative” matrix representing negative associations. This separation avoids the cancellation of opposite effects when aggregating the maps to coarser levels (see below).
2. Region level: We computed the row-wise average for each region in the “positive” and “negative” matrices. This yields two measures of how each region’s connectivity relates to the prediction. A region value can be high in both the positive and negative maps, as it can have strong positive associations for some connections and strong negative ones for others.
3. Network level: Each cortical region was assigned to one of 17 functional networks as defined in Yeo et al. (2011). The 19 noncortical regions were considered as a single additional network. Separating them by brain hemisphere yielded 36 functional networks. On this basis, between– and within-network feature importance was computed by averaging the corresponding Haufe-transformed weights for each network (within-network effect) and for each network pair (between-network effect), again separately for the negative and positive FC-level Haufe-transformed weights.

To benchmark these data-driven maps against more conventional approaches, we also computed mass-univariate Pearson correlations between each FC and the outcome variable, after deconfounding the latter for age, sex, and head motion. This approach is commonly referred to as a brain-wide association study (BWAS). Prior results from the UKBB exploratory analysis and recent literature (Chen et al., 2023) have shown that Haufe transform maps tend to be more reliable and generalizable than the linear associations highlighted by BWAS. Following the same aggregation strategy, we computed BWAS maps at the connection, region, and network levels.

### Statistical analysis

To test our hypotheses about model performances, we evaluated whether the MMM-KRR model outperforms the baseline KRR and whether MMM-EN provides further improvement. Specifically, we compared the distributions of 100 correlations obtained from repeated train-test splits for each model. These pairwise comparisons were assessed using one-tailed paired t-tests on the correlation distributions, with correction for repeated splits (Nadeau & Bengio, 2003), as the models were trained over the same subjects in each repetition.

We empirically assessed the significance of *r* > 0 for each model performance using permutation testing. Specifically, we repeated the analysis 1,000 times after randomly shuffling the outcome variables across subjects. For each permutation, the results were estimated as the average over 100 repetitions. This procedure generated null distributions for both the performance metrics and the relevance maps. Empirical p-values were computed as the proportion of permutations yielding an average performance equal to or greater than the observed model performance. To correct for the high number of elements in the relevance maps, we applied false discovery rate (FDR) correction (*q* = 0.05) for multiple comparisons within each map.

We assessed the split-half reliability of the relevance maps from the best-performing model by splitting the ReCoDe dataset into two independent halves, refitting the model, and generating them for each half. We assessed reliability as the pairwise Pearson’s correlation between the two maps for each level, averaged over 100 data splits. To investigate the generalizability of our relevance maps, we applied the same approach to compare the maps generated for the whole ReCoDe cohort with those from our literature review (available at https://osf.io/gs5b4 for download and at https://connexplore.med.tu-dresden.de for visualization), the UKBB subsample, as preregistered, and the BWAS maps.

For data-driven relevance maps to be meaningful, they must also be outcome-specific, so that different outcomes yield distinct maps. If the same connectivity patterns are highlighted regardless of the predicted variable, the maps would lack interpretability and biological relevance. To test this, we generated relevance maps for predicting age, a variable independent of AUDIT-C, since it is one of the confounders regressed out of AUDIT-C, and evaluated their similarity (Pearson correlation) with the AUDIT-C maps. The significance of the similarities between relevance maps was tested with a spin test, which controls for spatial autocorrelation by rotating cortical regions on an inflated surface to generate 10,000 null atlas configurations. These null configurations are then used to create 10,000 null-relevance matrices at the connection level and, subsequently, to generate a null distribution of similarities between the permuted importance maps. Statistical significance is determined as the proportion of null similarities that exceed the observed correlation. Since the spin test can only be applied to cortical regions, the 19 noncortical regions are excluded when calculating the p-value.

### Sensitivity analyses

As sensitivity analyses, we ran alternative analysis approaches, like including the number of DSM-5 criteria for TUD as an additional confounder and, separately, excluding participants with framewise displacement (FD) > 0.5 mm in more than 7.5% of frames (an additional N=104 participants excluded), to assess the robustness of model performance to these factors. In addition, we repeated the analyses in subsamples of N=300 participants to compare the KRR performance across sex, age, and AUD severity groups. Specifically, we compared males and females; younger (≤30 years) and older (>35 years) participants; and participants with fewer than 4 DSM-5 AUD criteria (no or mild AUD) versus those with 4 or more criteria (moderate-to-severe AUD).

## Results

Table 1 summarizes the main demographics and clinical characteristics of the cohorts used in this analysis. The two datasets were matched for gender and drinking patterns, as described in the preregistration. The cohorts differ substantially in age, with ReCoDe being significantly younger (mean = 35.32, range = 18–65) compared to the UKBB sample (mean = 63.01, range = 40–81). The UKBB also shows a significantly lower mean FD, indicating less head motion during scanning, but a higher AUDIT-C score, likely due to differences in the definition of a standard drink between the UK and Germany. Quantity-Frequency surveys and Structured Clinical Interviews for DSM-5 (SCID) were conducted only in the ReCoDe sample, revealing that 528 participants (71.4%) met the definition for AUD and 225 (30.4%) for tobacco use disorder (TUD) (more than 1 criterion); 156 individuals (21%) met criteria for both AUD and TUD.

**Table 1:**
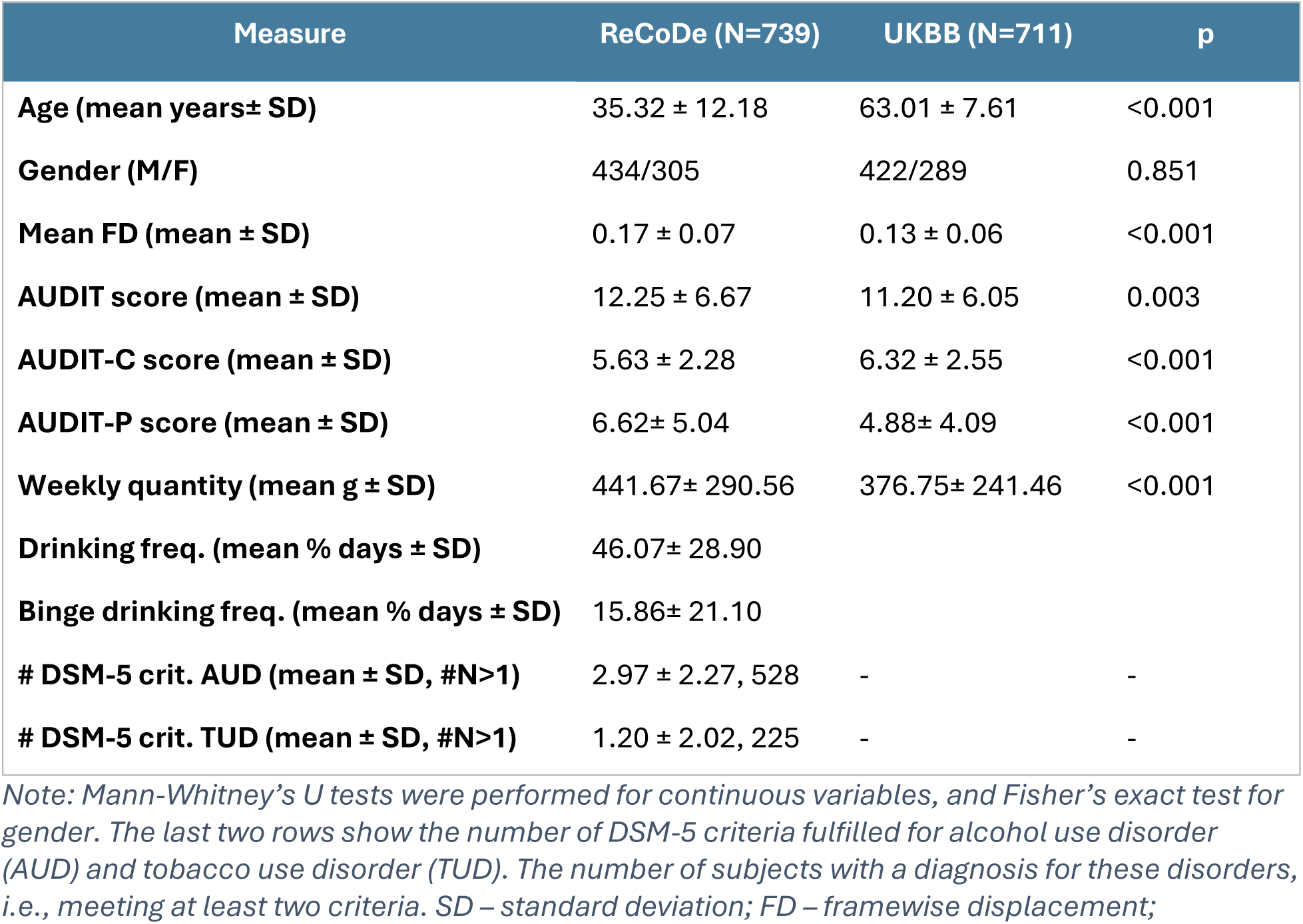
Baseline measures for the datasets employed.

As illustrated in Figure 1, the models based on the MMM embedding did not outperform the locally trained KRR models, unlike what we observed in the original UKBB subsample.

**Figure 1:**
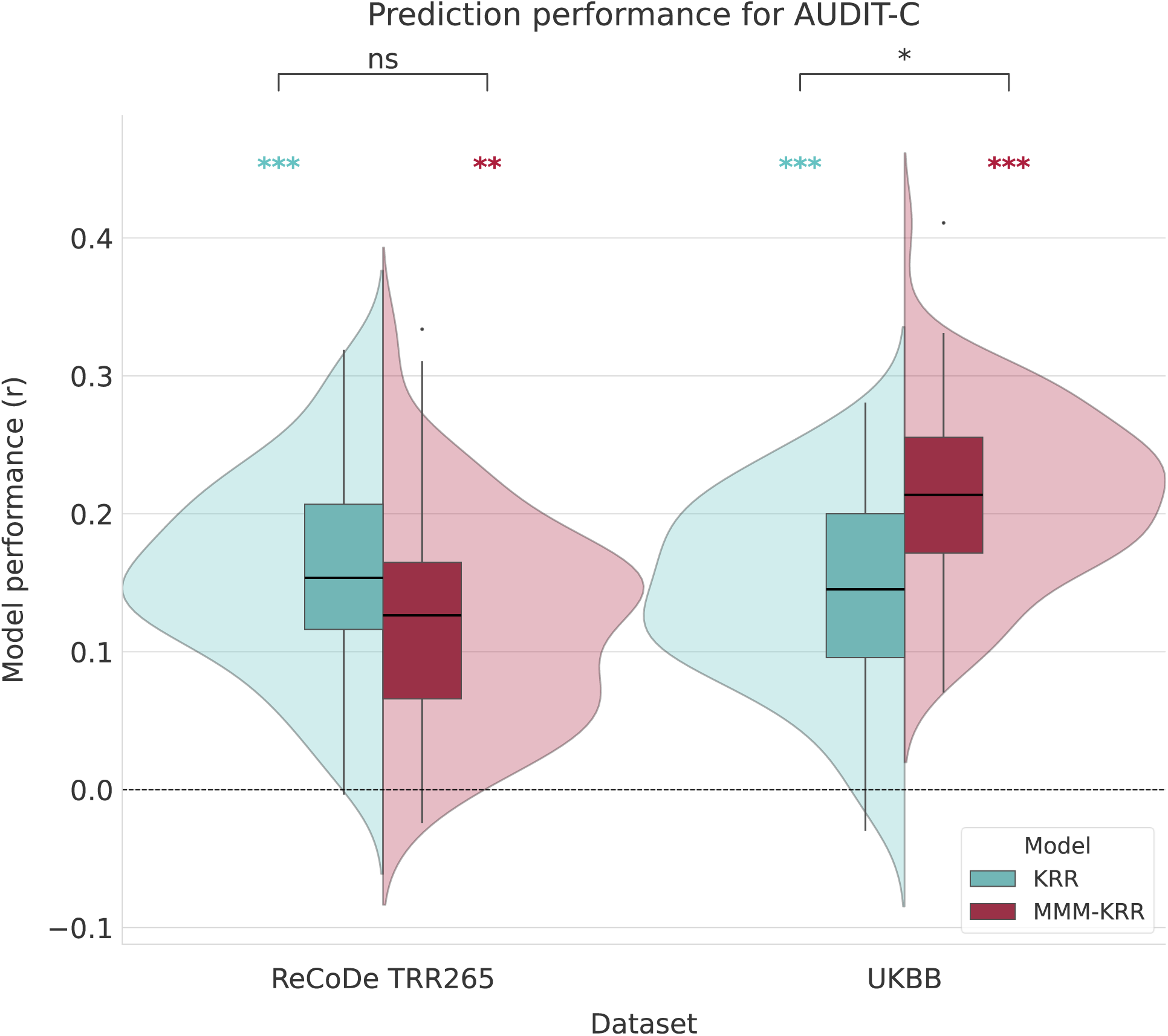
Performances of the models predicting AUDIT-C score investigated in ReCoDe dataset (N=73S) and in the UK Biobank subsample (UKBB, N=711). Violin and box plots show the distribution of test-set correlations across 100 train–test splits. Performance differences between models were assessed using corrected one-tailed paired t-tests (on top). Significance of individual models (shown above each violin) was estimated from a permutation-based null distribution. Significance levels are expressed as *: p <.05, **: p<.01, ***: p<.001

Both KRR and MMM-KRR show highly significant performance, with the average *r* = 0.15. In our preregistered findings on UKBB, the MMM-EN model significantly outperformed both KRR and MMM-KRR. In the ReCoDe cohort, however, the model failed to converge on most splits, proving unstable when performance was lower. The results for all other outcomes, including age, are reported in Figures S1 and S2 in the supplementary materials. Overall, for no outcome, including age, MMM led to a significant performance improvement, and the AUDIT score and its subscales performed, on average, better than the other AUD-related outcomes. As for the sensitivity analyses, excluding subjects with the highest head motion in the scanner (N = 104) did not significantly affect model performance, whereas regressing out the number of TUD criteria from the outcome yielded slightly worse performance, as shown in Figure S3 of the supplementary materials. When repeating the analyses in more homogeneous subsamples, prediction performance was significantly higher in younger than in older participants, and in participants with no or mild AUD than in those with moderate-to-severe AUD, as shown in Figure S4 in the supplementary materials.

We show the relevance maps for predicting AUDIT-C with MMM-KRR in Figure 2, highlighting positive and negative associations at the regional and network levels, respectively. A visualization of the relevance map at functional connectivity level can be found at https://connexplore.med.tu-dresden.de.

**Figure 2:**
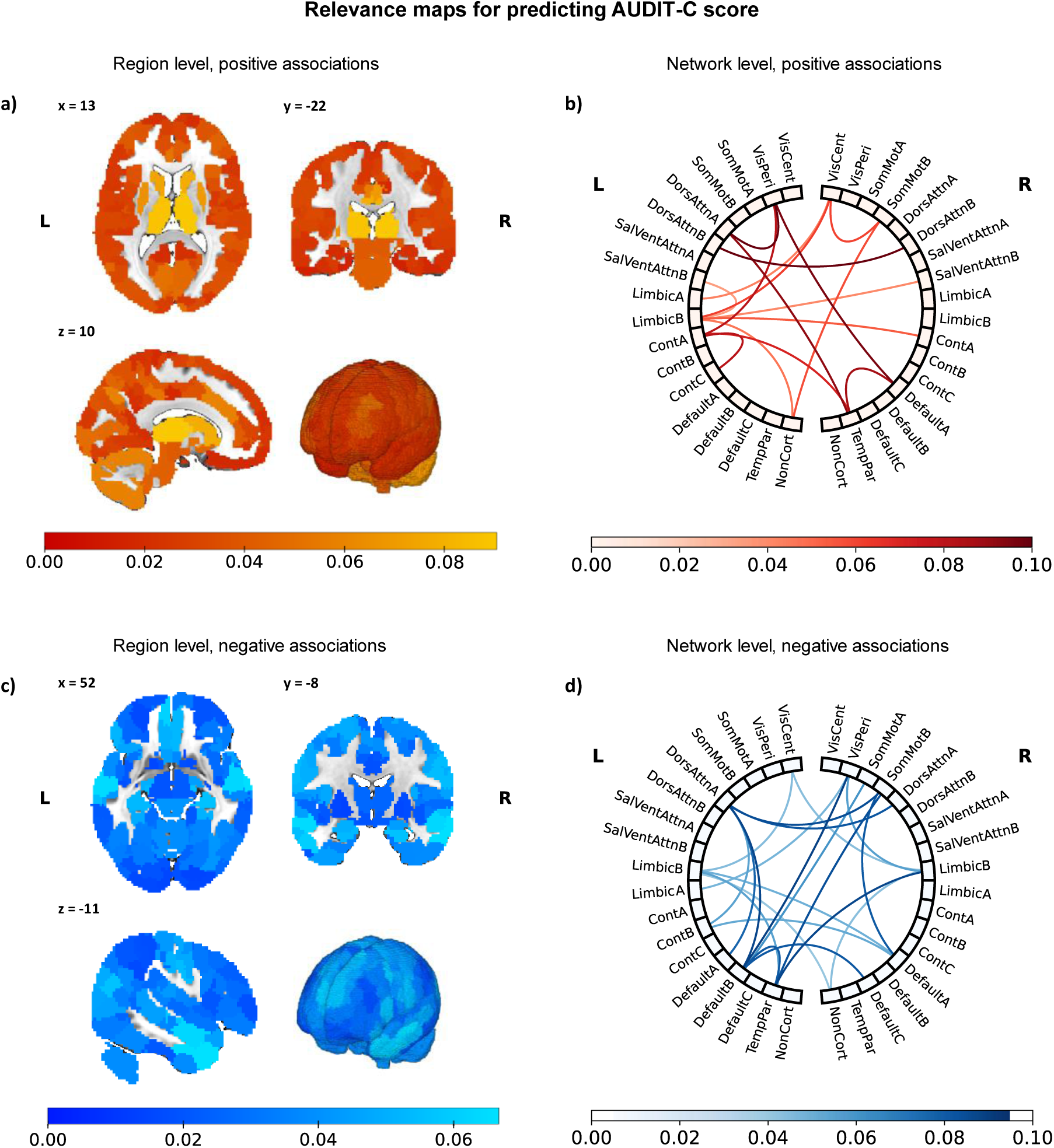
Relevance maps for AUDIT-C prediction from the MMM-KRR model at the region and network levels. (a,c) Region-level relevance for positive (a) and negative (c) associations, mapped onto anatomical space with Schaefer+ atlas. Colors indicate the strength of the association between altered functional connectivity and predicted AUDIT-C scores (absolute correlation). Axial (top-left), coronal (top-right), sagittal (bottom-left), and 3D rendering (bottom-right) views are shown in MNI space. (b,d) Network-level relevance for positive (b) and negative (d) associations. Chords indicate between-network associations that were significant according to spin tests (p_spin_ < 0.05, uncorrected). Node colors, on the other hand, reflect within-network relevance.

Regarding the positive associations of the predicted AUDIT-C score with regional connectivity, the highest relevance values (Figure 2a) are observed in the bilateral posterior cingulate cortex (PCC), the precuneus, and particularly in the bilateral thalamus and dorsal striatal regions, such as the putamen and caudate. At the network level (b), the chord diagram highlights higher connectivity in a circuit formed between the right DMN-A (precuneus, PCC, frontopolar cortex), right temporoparietal network (superior temporal gyrus), the bilateral dorsal attention network A (DAN–A; superior parietal lobe), and the left visual peripheral network (angular gyrus, cuneus), as well as bigger overall centrality of the left limbic network B (orbitofrontal cortex).

On the other hand, the most negative associations at the regional level are observed in the bilateral middle temporal gyrus (MTG), the dorsomedial prefrontal cortex (dmPFC), the left inferior frontal gyrus (IFG), and limbic structures, including the hippocampus and amygdala (Figure 2c). At the network level (Figure 2d), the chord diagram highlights reduced connectivity between the left DMN-B (IFG, anterior temporal pole, dmPFC), the right sensorimotor network (SMN-B; granular insula, opercular postcentral gyrus), the bilateral DAN-A, and the bilateral limbic network B.

The relevance maps derived from the KRR model showed more modest associations, which largely failed to reach significance in the spin test, yet remained highly correlated with the corresponding MMM-KRR maps (*r* ≈ 0.7), as shown in Table S1 in the supplementary materials. The full relevance maps are provided in the supplementary materials.

In Figure 3, we show the similarity between the relevance maps for two independent outcomes, derived in the ReCoDe cohort, and their counterparts derived from the preregistered UKBB analysis and from our previous literature review. For AUDIT-C, Haufe-transformed maps show moderate but significant cross-cohort reliability at the FC level and for negative associations at the region– and network-levels. These effects are consistently stronger than in BWAS maps. MMM-KRR yields slightly higher cross-cohort similarity overall compared to KRR, but at the cost of reduced specificity, as reflected in increased cross-outcome similarity (e.g., up to *r* = 0.38), whereas KRR maps remain largely outcome-specific. In contrast, age-related maps exhibit uniformly lower cross-cohort similarity. These findings suggest a trade-off between reliability and outcome specificity, with MMM-KRR producing more stable but less outcome-specific relevance patterns. Finally, correspondence with maps derived from the literature review is negligible, with only a small effect for the KRR AUDIT-C negative region-level map (r = 0.13), which is significant after the spin test.

**Figure 3:**
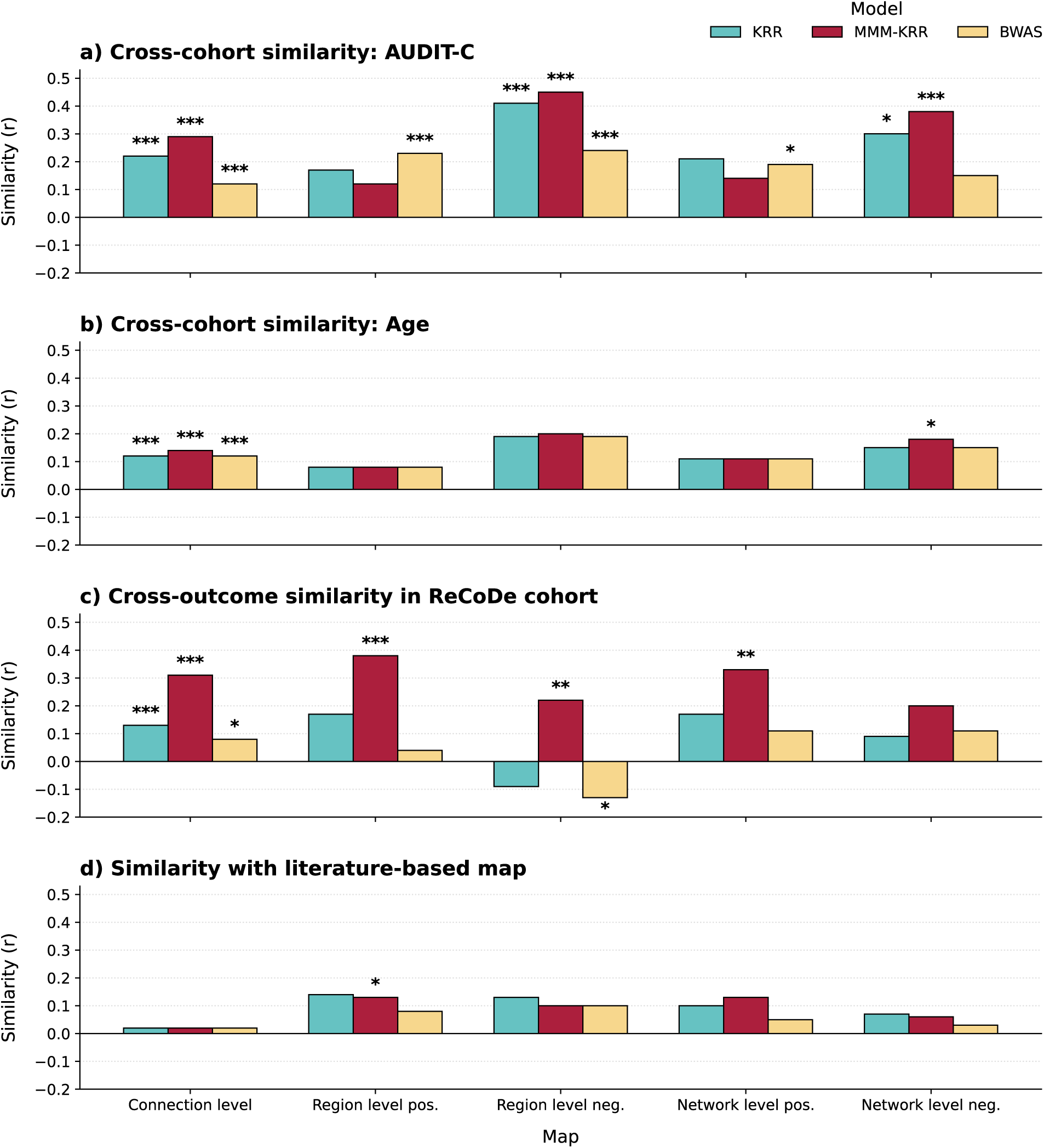
Pearson correlation similarity between maps derived using KRR, MMM-KRR, and BWAS at the connection, region, and network levels. Panels show cross-cohort similarity between the UK Biobank subsample and the ReCoDe cohort for (a) AUDIT-C and (b) age; (c) cross-outcome similarity between AUDIT-C and age maps within the ReCoDe cohort; and (d) similarity between the ReCoDe AUDIT-C maps and the literature-derived maps from the systematic review. Region– and network-level results are shown separately for positive and negative associations. Significance is tested with spin-tests, with significance levels expressed as *: p <.05, **: p<.01, ***: p<.001

Split-half reliability is highly dependent on the specific subject split for small effect sizes, with substantial variability across splits. Across all models, AUDIT-C showed generally low split-half reliability, although moderate effect sizes were observed at the region level for MMM-KRR. In contrast, age relevance maps show consistently high and significant split-half reliability across all models and levels of aggregation. Further details on this analysis are reported in the supplementary materials (Figure S4).

## Discussion

In this preregistered study, we investigated the association between rs-fMRI FC matrices and alcohol consumption in a large cohort of non-treatment-seeking AUD subjects and controls (ReCoDe). We employed machine learning algorithms and specifically considered the potential benefit of meta-matching transfer learning. We compared the predictive performance of locally trained regression models with that of meta-matching models. We interpreted the models by extracting data-driven relevance maps using the Haufe transform to evaluate the linear contribution of each input feature, and by assessing split-half reliability, outcome-specificity (AUDIT-C versus age), and generalizability. The latter was assessed across cohorts, using preregistered maps from the UK Biobank (10.17605/OSF.IO/SJZY9) and across the literature based on our previous review (Bottino et al., 2025).

We found that machine learning models can predict AUDIT-C scores with *r* ≈ 0.15. However, we could not replicate the performance increase observed in our preregistration and in previous literature (Chen et al., 2024; Chopra et al., 2024) through meta-matching. One possible explanation for the decline in the effectiveness of the meta-matching models is partial data overlap between the UKBB exploration sample and the data used to train them, i.e., a form of data leakage that might have inflated the performance reported in the exploratory findings. In addition, the meta-matching framework has been shown to provide an edge in prediction when applied to datasets up to 200 subjects (He et al., 2022), whereas ReCoDe comprised 739 participants. At this sample size, training directly on the target phenotype may recover cohort-specific information that is lost when predictions rely on a lower-dimensional representation learned from external datasets. Consistent with this interpretation, meta-matching also provided no advantage for age, with prediction substantially stronger in ReCoDe than in UKBB (Figure S2), likely reflecting the broader age range of the ReCoDe cohort.

Moreover, in a recent study by Hearne et al. (2026), meta-matching did not outperform a locally trained logistic regression model in distinguishing individuals with obsessive compulsive disorder from healthy controls, suggesting, together with our findings, that the benefits of meta-matching may be less pronounced for psychiatric phenotypes than for cognitive outcomes.

Overall, AUDIT-C prediction from FC remains modest, which may be due to several factors. Drinking patterns have previously shown poor predictability in big datasets, as shown, for example, in He et al. (2022), with performance for predicting alcohol intake around *r* ≈ 0.1, in line with previous results on behavioral prediction of non-cognitive measures, which have shown less predictability than cognitive ones in a range of studies (Chen et al., 2022; Marek et al., 2022; Ooi et al., 2022). Moreover, the ReCoDe dataset comprises mostly mild, non-treatment-seeking subjects, with an average of 3 DSM-5 criteria out of 11, as shown in Table 1. While this cohort might be more representative of moderate drinkers than of severe clinical cases, it shows limited outcome variance, which may contribute to the low general predictive performance across all AUD-related outcomes (see Figure S1 in the supplementary materials). Finally, resting-state FC might not be altered by current drinking patterns to the same extent as by lifetime exposure. For example, Weiland et al. (2014) found that the association between years of regular drinking and AUD severity was mediated by reduced ECN connectivity.

To gain scientific insights into brain-behavior relations, we studied data-driven feature-relevance maps. We compared the reliability of maps for two independent outcomes (age and AUDIT-C score) across different cohorts and across two halves of the ReCoDe cohort, for three map types: Haufe maps derived from the KRR model, the ones from the MMM-KRR models, and univariate correlation maps (BWAS), with Haufe-transform-based maps being, on average, more reliable than BWAS maps. Our findings are in line with this for targets with low predictability.

The biological interpretation of the relevance maps aligns with previous results in the literature and with our hypothesis, particularly regarding the negative associations at the regional and network levels, which were replicated across cohorts (Figure 3). Specifically, higher predicted alcohol consumption was associated with reduced connectivity involving the left inferior frontal gyrus (IFG), a region strongly implicated in inhibitory control (Su et al., 2025), the dorsomedial prefrontal cortex (dmPFC), and, most prominently, anterior temporal regions. Reduced connectivity between regions of the limbic network (amygdala, OFC, and anterior MTG) with the DAN and the dmPFC suggests alterations in emotion and reward processing with potentially reduced top-down attentional control (Zhang & Volkow, 2019). We also observed patterns consistent with hypothesized hyperconnectivity associated with alcohol consumption, including increased connectivity between regions of the striatal reward system, such as the caudate and putamen, DMN parietal regions, such as the PCC and precuneus, and the parietal DAN. Notably, several of these regions have previously shown greater responses to alcohol cues in individuals with AUD than in healthy controls (Schacht et al., 2013), suggesting a possible link with enhanced processing of alcohol-related motivational or contextual information. However, these positive associations did not reach statistical significance and showed more limited reliability, and should therefore be interpreted cautiously. Overall, the replicated findings primarily point to alterations in the DMN, particularly its interactions with temporal and limbic regions, suggesting a more central role for DMN-related alterations than for the broader triple-network model in alcohol consumption.

This study has several limitations. First, its cross-sectional design precludes inferences about causal relationships between FC patterns and alcohol use, making it difficult to determine whether the observed associations reflect consequences of prolonged alcohol exposure or pre-existing neural characteristics that predispose individuals to higher alcohol consumption. Future longitudinal studies could help disentangle these mechanisms by tracking changes in both alcohol use and brain connectivity over time and by examining their relationship with structural markers of alcohol-related brain alterations, such as gray matter volume loss or cortical thinning (Durazzo et al., 2015). Second, although the sample was relatively large, it consisted primarily of non-treatment-seeking individuals with mostly mild-to-moderate AUD severity and substantial demographic heterogeneity, which may have contributed to smaller effect sizes and, consequently, lower predictive performance and reliability of the derived relevance maps. Supporting this possibility, our exploratory subsample analyses showed significantly higher predictive performance in younger participants (≤30 years) compared with older participants (>35 years), and in participants with no-to-mild AUD compared with those with moderate-to-severe AUD. These findings suggest that demographic and clinical heterogeneity may partly obscure FC– alcohol consumption associations and highlight the potential value of studying more homogeneous populations. Future studies could extend this approach by identifying behaviorally homogeneous AUD subtypes and testing whether they are associated with more distinct and reproducible connectivity patterns.

## Conclusion

Our findings show that resting-state FC contains a small but reproducible signal associated with alcohol consumption, although its utility for individual-level prediction remains limited. Meta-matching did not improve prediction over models trained directly in the target cohort, suggesting that its benefits may be limited for psychiatric phenotypes at moderate sample sizes. Nevertheless, the partial replication of feature-relevance patterns across independent cohorts, particularly for reduced connectivity involving default mode, limbic, and dorsal attention networks, supports the potential of predictive models to identify reproducible brain-behavior associations even when predictive accuracy is modest.

## Acknowledgments

We thank the non-author collaborators of the ReCoDe Consortium, whose full names and affiliations can be found at https://www.trr265.org/en. Members of the consortium may not be in a position to endorse the content of the manuscript.

## Authors contributions

Conceptualization: MB, MM, MNS, HW

Data curation: MB, MM, MNS, PB, FK, BL, AH, GS, AR

Formal analysis: MB, MM

Funding acquisition: MM, HW, ReCoDe Consortium

Methodology: MB, MM, BTTY, JB, HW, KR, SG, RPR, ES

Writing – original draft: MB, JB, MM, MNS, BTTY, HW

Writing – review C editing: All authors

## Data and materials availability

The underlying MRI data and individual-level study data are not publicly available due to ethical and privacy restrictions. The relevance maps mentioned in the manuscript can be found in the supplements and visualized at https://connexplore.med.tu-dresden.de/. The scripts used for the analyses are publicly available on <u>OSF</u>.

## Ethics approval statement

The study was approved by the local ethics committees at Charité–Universitätsmedizin Berlin, Technical University Dresden, and the Central Institute of Mental Health Mannheim, under the application numbers EK 459112018 and SR-EK-165042023”.

## Participant consent statement

All participants provided written informed consent. For participants younger than 18 years, informed consent was co-signed by a parent or legal guardian.

## AI disclosure statement

Generative AI tools, including various versions of ChatGPT, were used to improve the manuscript’s clarity and readability and to assist in developing some of the analysis scripts. All AI-assisted content and code were reviewed and validated by the authors, who take full responsibility for the accuracy of the final manuscript.

