## Supplementary materials S1 for "Predicting alcohol consumption from resting-state functional connectivity: Feature relevance and the potential of meta-matching"

#### Title

*shared senior authorship

### Supplementary details on methods and results

#### Datasets matching

As explained in the preregistration, we decided to conduct an exploratory analysis using a subset of participants from the UK BioBank cohort (UKBB) who matched part of the ReCoDe cohort on drinking patterns and gender. UK Biobank is a large-scale biomedical database and research resource containing genetic, lifestyle, and health information from half a million UK participants. UK Biobank’s database, which includes blood samples, heart and brain scans, and genetic data of the 500.000 volunteer participants, is globally accessible to approved researchers who are undertaking health-related research that’s in the public interest.

To build the matching subset, we applied a nearest-neighbor matching approach: for each subject in the ReCoDe cohort, we identified the closest UKBB participant of the same sex who had not already been matched to another ReCoDe subject. The similarity metric was the Euclidean distance computed from a set of drinking-related variables after standardizing them to have zero mean and unit variance.

We visually inspected the distributions of the drinking patterns in this new dataset of 712 subjects to be similar to those in the ReCoDe cohort. Matching the age distribution was not possible as the two cohorts cover different age ranges match (ReCoDe: 16-65 years old; UKBB: 45-82 years old) that would be impossible to match.

#### Imaging details

Scanning was carried out on 3T MRI scanners (Siemens Magnetom Prisma and Trio) using the Siemens 64-channel head coil in Berlin and Mannheim (Prisma systems) and the 32-channel head coil in Dresden (Trio System and Prisma Fit starting March 2022).
Functional images were acquired using a multi-band T2*-weighted gradient echo-planar imaging (EPI) sequence (500 volumes, TR = 869 ms, TE = 38 ms, 60 slices, slice thickness = 2.4 mm, voxel size 2.4 × 2.4 × 2.4 mm, no inter-slice gap, field of view (FoV) = 210 mm, matrix size 88 x 88, acquisition orientation T > C, interleaved slice order, multi-band factor 6, flip angle = 58°, bandwidth = 1832 Hz/Px, prescan normalize, weak raw data filter, fat sat).
Field map images for distortion correction were obtained using a Siemens dual gradient-echo sequence (TR = 698 ms, TE1 = 5.19 ms, TE2 = 7.65 ms, 64 slices, slice thickness = 2.4 mm, no slice gap, voxel size 2.4 × 2.4 × 2.4 mm, field of view (FoV) = 210 mm, matrix size 88 x 88, acquisition orientation T > C, interleaved slice order, flip angle = 54°, bandwidth = 279 Hz/Px).
T1-weighted MPRAGE images were used for normalization (TR = 2000 ms, TE = 2.01 ms, TI = 880 ms, FoV = 256 mm, 208 sagittal slices, voxel size 1 × 1 × 1 mm, flip angle = 8°, GRAPPA factor 2 (PE), lines, prescan normalize, 23.1% slice oversampling, bandwidth = 240 Hz/Px).

#### Machine learning models

KRR is a variant of Ridge regression that operates in a kernel-defined feature space, where the kernel function measures similarity between the training subjects. A common approach in the literature for regression of behavioral outcomes from FC is to use Pearson’s correlation between the input arrays as a measure of similarity in the kernel space (He et al., 2020). The only hyperparameter is the regularization strength of the Ridge component, searched logarithmically in the range ${10}^{-2}-{10}^{4}$

ElasticNet is a linear regression model that combines L1 (Lasso) and L2 (Ridge) regularization to control overfitting. The model hyperparameters are the regularization strength α, searched logarithmically in the range ${10}^{-1}-{10}^{3}$ and the ratio between L1 and L2 penalties, searched linearly in the range $0.1-0.9$.

#### Discussion over split half reliability

The split-half reliability of these maps varied substantially across the 100 random half-splits (Figure S5), indicating that feature importance estimates were highly dependent on sample composition. Although we observed the same trends reported by Chen et al. (2023), for which Haufe-derived relevance maps showed higher reliability than BWAS maps, we did not observe significant reliability for AUDIT-C. One possible explanation is the relatively low signal-to-noise ratio of addiction-related phenotypes, which likely limits both predictive performance and the stability of feature importance estimates. Consistent with this interpretation, age showed moderate-to-high reliability across all models, reflecting its stronger association with FC and higher prediction accuracy. This supports the conclusion of Chen et al. (2023) that feature importance reliability is closely linked to model performance. A second explanation is limited sample size: Chen et al. (2023) have already shown in their work that split-half reliability for sample sizes under 1000 subjects is generally very low. While larger samples could improve the stability of relevance maps, using more demographically homogeneous cohorts, e.g., focusing on subgroups of AUD, could reduce the need for aggressive deconfounding procedures that, in our case, account only for linear differences between samples. Finally, the strong collinearity of FC features may further reduce the stability of feature importance estimates. Notably, although meta-matching did not improve predictive performance in our dataset, it produced more reliable relevance maps, potentially because the pretrained embeddings reduce redundancy in the input feature space. However, this increase in reliability was accompanied by greater similarity between relevance maps of different outcomes, suggesting that dimensionality reduction may improve stability at the cost of outcome specificity due to information loss during the embedding.

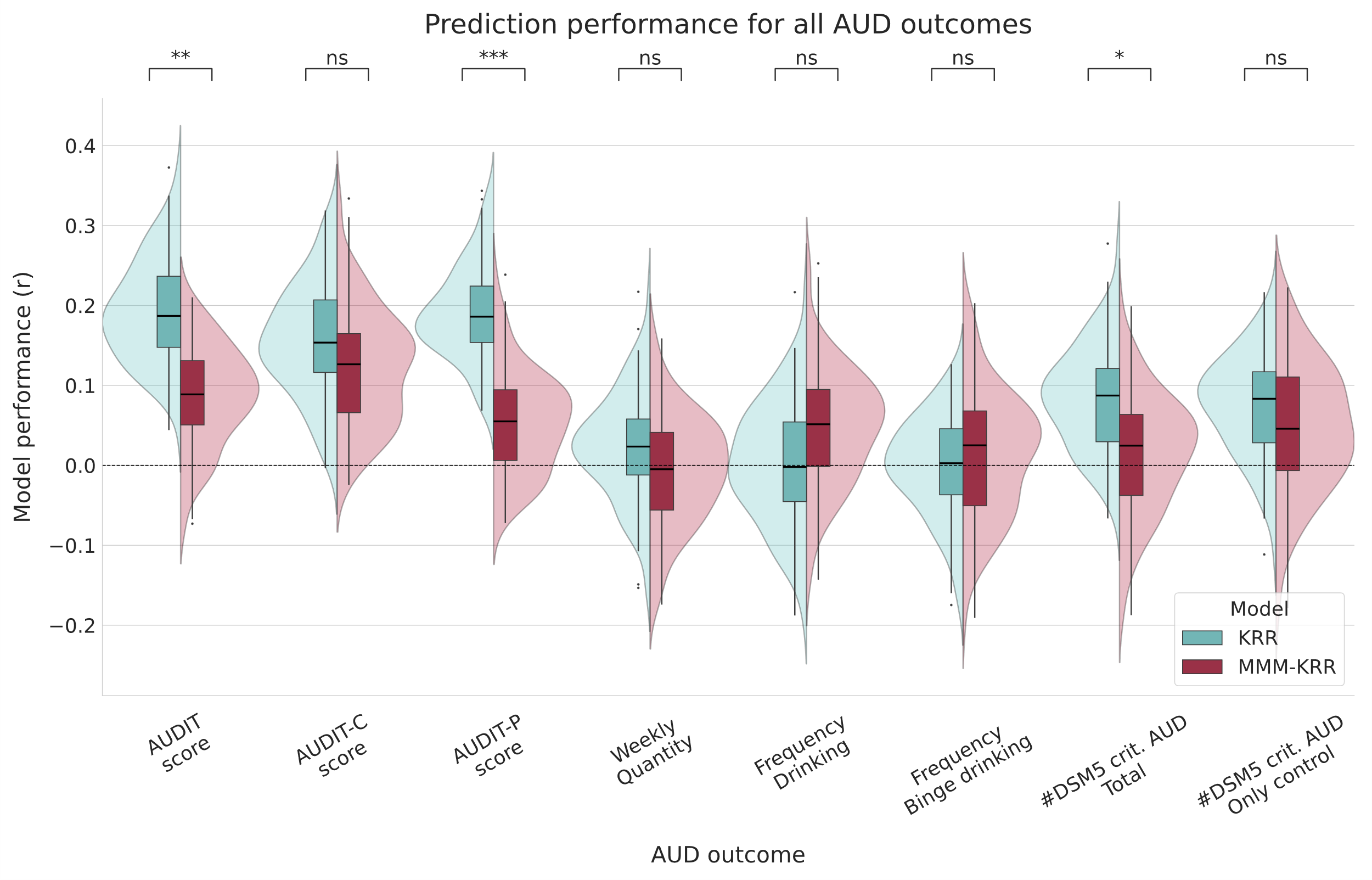

Figure S1: Performances of the models predicting alcohol use disorder (AUD)-related outcomes in the ReCoDe cohort (N=739). Violin and box plots show the distribution of test-set correlations across 100 train–test splits. Performance differences between models were assessed using corrected one-tailed paired t-tests (on top). Significance levels are expressed as ns: p >.05, *: p <.05, **: p<.01, ***: p<.001

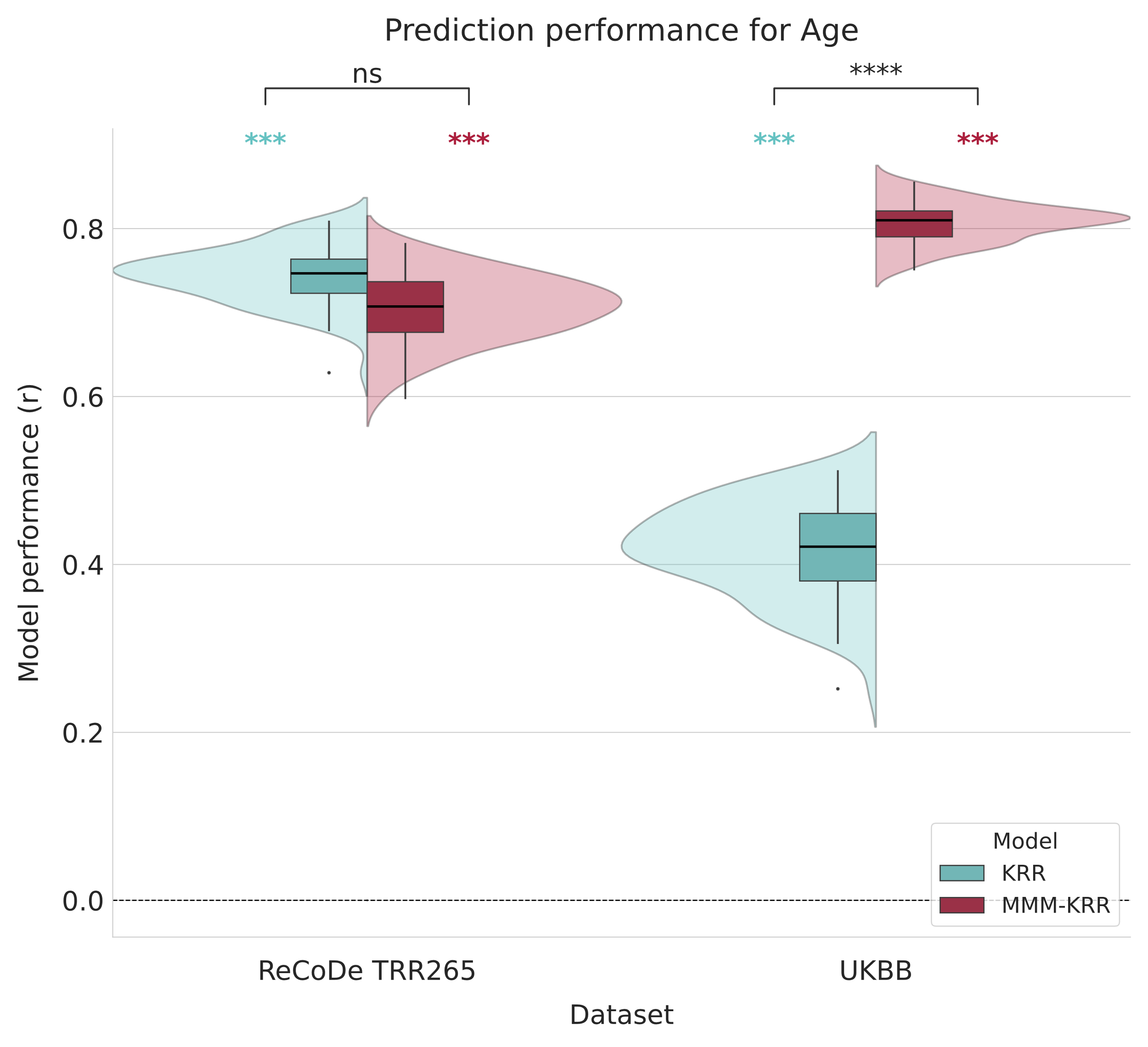

Figure S2: Performances of the models predicting participants’ age investigated in the ReCoDe cohort (N=739) and in the UK Biobank subsample (UKBB, N=711). Violin and box plots show the distribution of test-set correlations across 100 train–test splits. Performance differences between models were assessed using corrected one-tailed paired t-tests (on top). Significance of individual models (shown above each violin) was estimated from a permutation-based null distribution. Significance levels are expressed as ns: p >.05, *: p <.05, **: p<.01, ***: p<.001

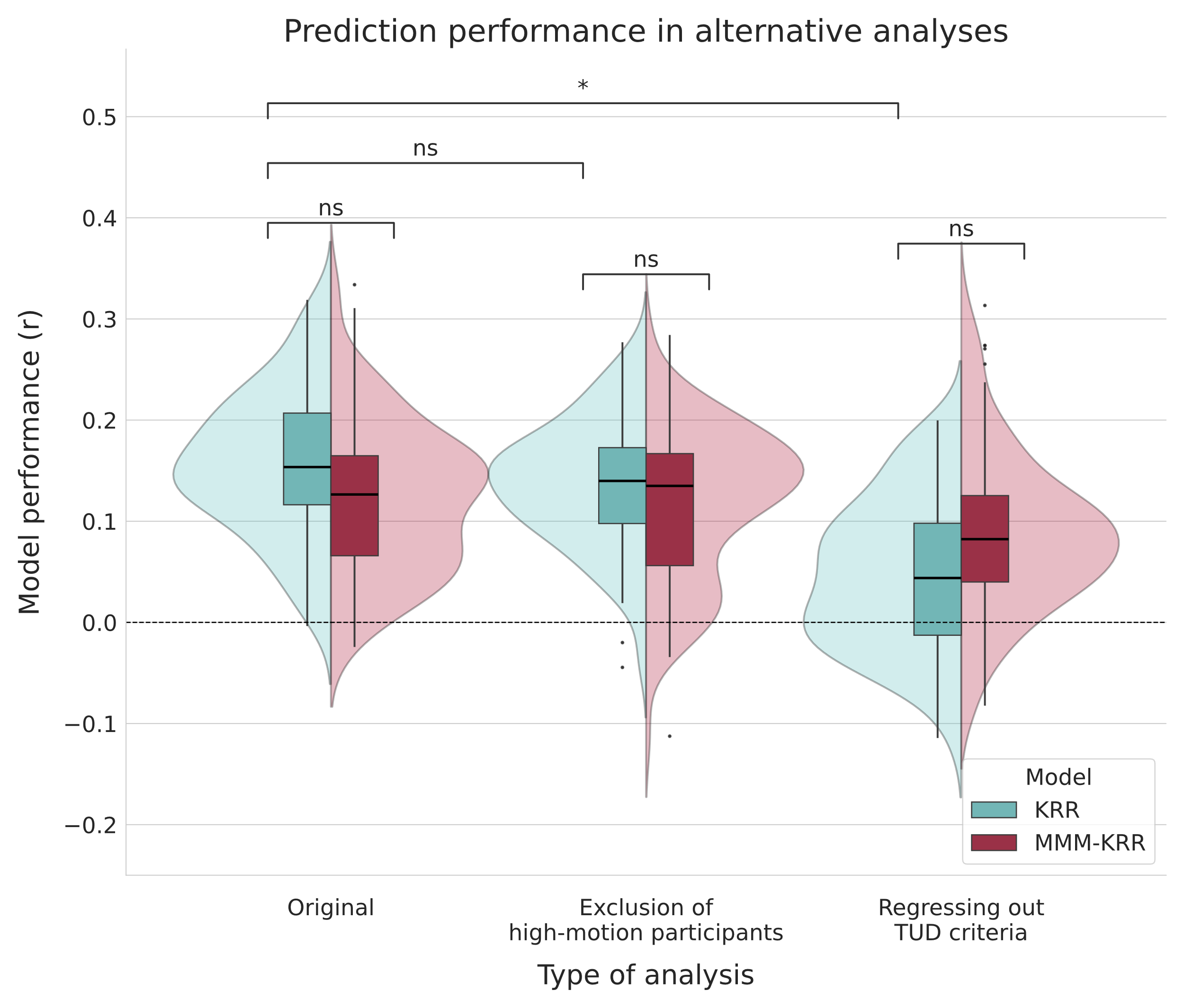

Figure S3: Performances of the models predicting AUDIT-C score investigated in the ReCoDe cohort (N=739), for different types of analysis. Violin and box plots show the distribution of test-set correlations across 100 train–test splits. Performance differences between models were assessed using corrected one-tailed paired t-tests (on top). Significance levels are expressed as ns: p >.05, *: p <.05, **: p<.01, ***: p<.001

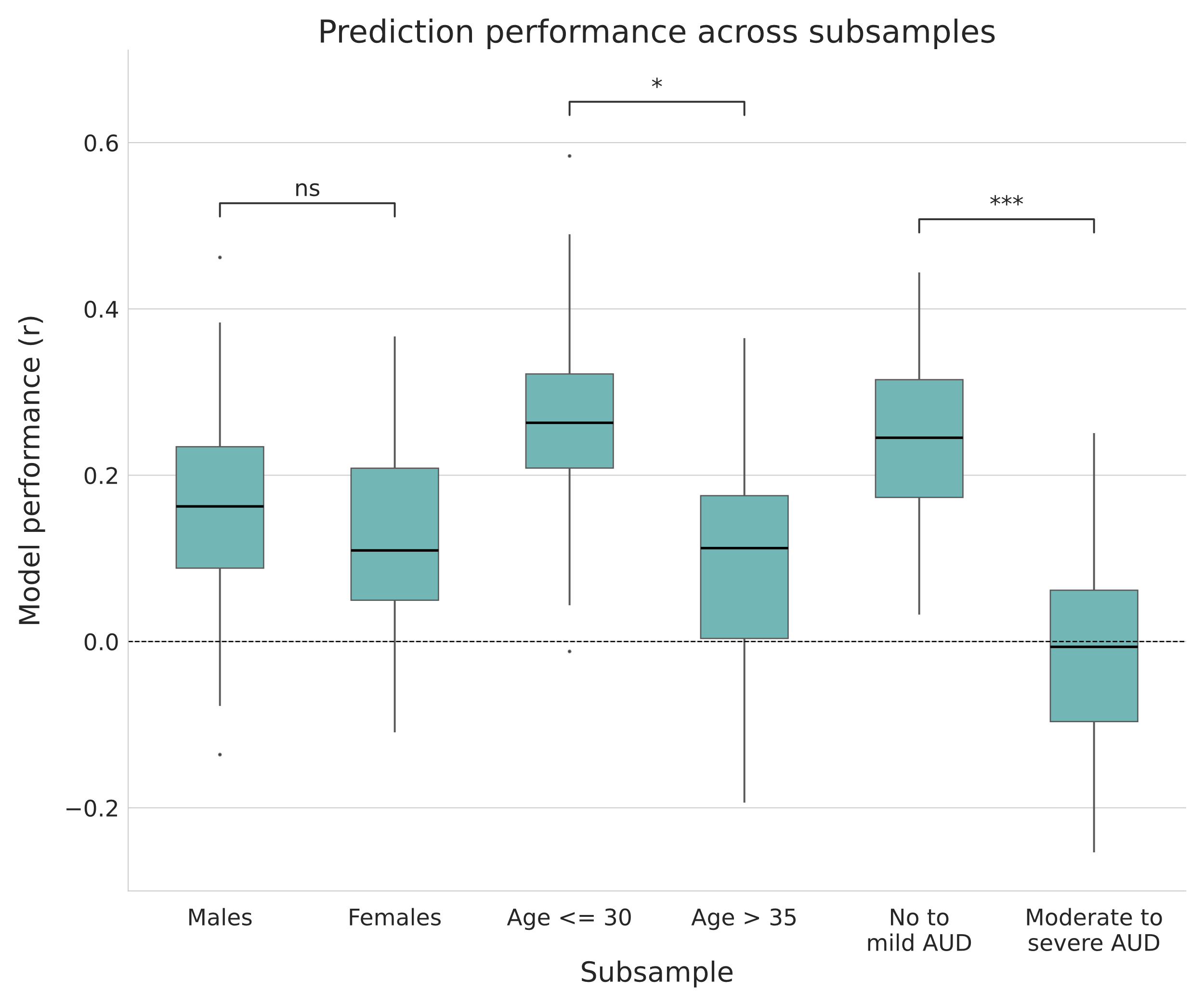

Figure S4: Performances of the KRR models predicting AUDIT-C score investigated in subsamples of the ReCoDe cohort (N=300). The “No to mild AUD” subsample only includes participants meeting 3 or less criteria for alcohol use disorder criteria according to the DSM-V, while the “Moderate to severe AUD” only includes participants meeting more than 3 criteria. The box plots show the distribution of test-set correlations across 100 train–test splits. Performance differences between subsamples were assessed using corrected two-tailed t-tests (on top). Significance levels are expressed as ns: p >.05, *: p <.05, **: p<.01, ***: p<.001

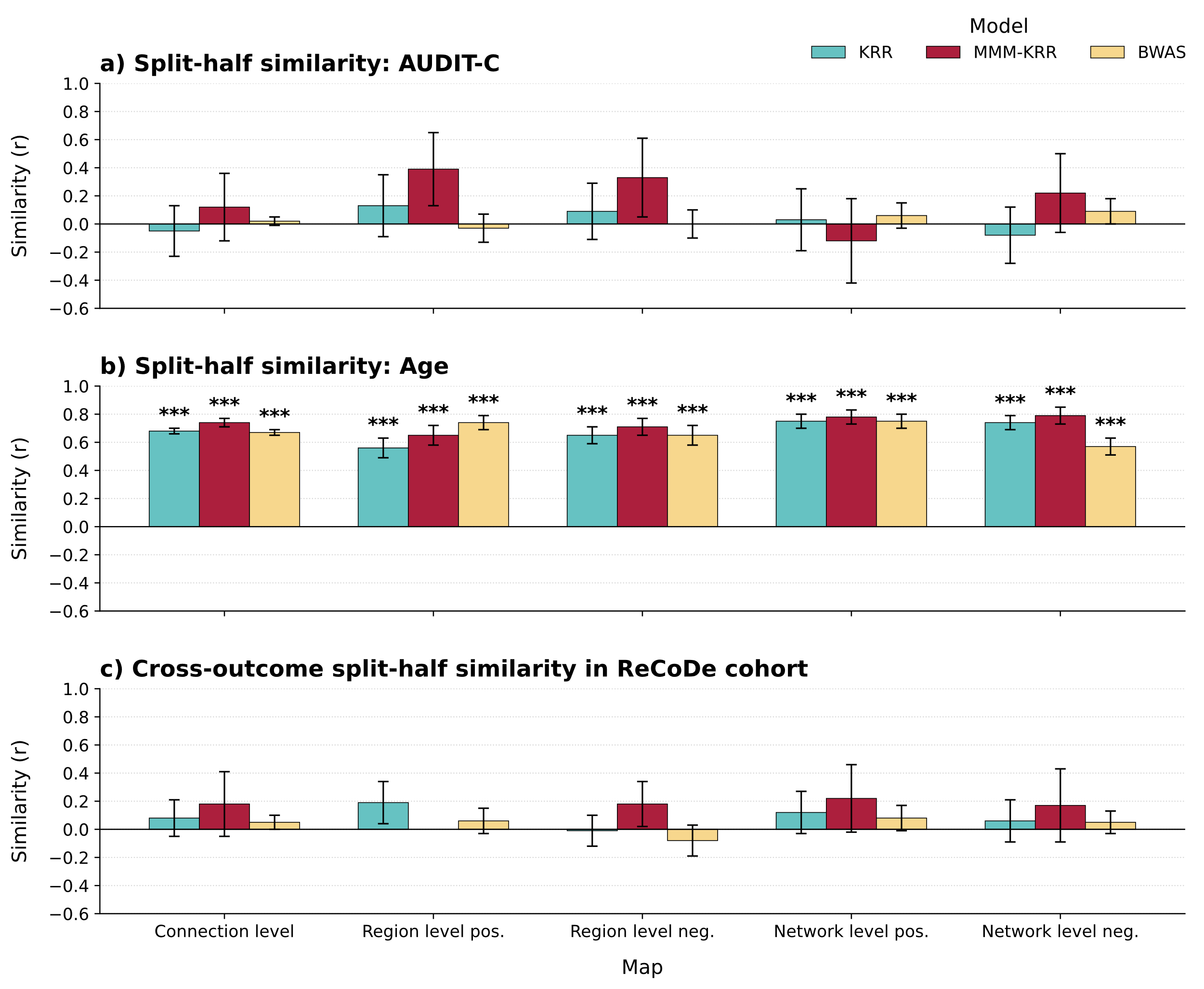

Figure S5: Average Pearson correlation similarity between maps derived using KRR, MMM-KRR, and BWAS at the connection, region, and network levels. Panels show split-half similarity in the ReCoDe cohort for (a) AUDIT-C, (b) age, and (c) cross-outcome split-half similarity between AUDIT-C and age maps. Region- and network-level results are shown separately for positive and negative associations. Error bars show one standard deviation of the similiarities over the 100 splits. Significance is tested with a corrected t-test against zero, with significance levels expressed as *: p <.05, **: p<.01, ***: p<.001

Table S1: Similarities between relevance maps for AUDIT-C score across models

| Map | Similarity  KRR / MMM-KRR | Similarity  KRR / BWAS | Similarity  MMM- KRR / BWAS |
| --- | --- | --- | --- |
| Connection level | 0.65 | 0.91 | 0.44 |
| Region level, pos. | 0.73 | 0.93 | 0.52 |
| Region level, neg. | 0.71 | 0.90 | 0.50 |
| Network level, pos. | 0.69 | 0.89 | 0.45 |
| Network level, neg. | 0.61 | 0.86 | 0.30 |
